# Two haplotype-resolved genomes of highly heterozygous AAB allotriploid bananas provide insights into subgenome asymmetric evolution and banana wilt control

**DOI:** 10.1101/2023.03.28.534356

**Authors:** Wen-Zhao Xie, Yu-Yu Zheng, Weidi He, Fangcheng Bi, Yaoyao Li, Tongxin Dou, Run Zhou, Yi-Xiong Guo, Guiming Deng, Wen-Hui Zhang, Min-Hui Yuan, Pablo Sanz-Jimenez, Xi-Tong Zhu, Xin-Dong Xu, Zu-Wen Zhou, Zhi-Wei Zhou, Jia-Wu Feng, Siwen Liu, Chunyu Li, Qiaosong Yang, Chunhua Hu, Huijun Gao, Tao Dong, Jiangbo Dang, Qigao Guo, Wenguo Cai, Jianwei Zhang, Ganjun Yi, Jia-Ming Song, Ou Sheng, Ling-Ling Chen

**Affiliations:** Institute of Fruit Tree Research, Guangdong Academy of Agricultural Sciences; Key Laboratory of South Subtropical Fruit Biology and Genetic Resource Utilization, Ministry of Agriculture and Rural Affairs; Guangdong Provincial Key Laboratory of Tropical and Subtropical Fruit Tree Research, Guangzhou 510640, China; College of Informatics, Huazhong Agricultural University, Wuhan 430070, China; State Key Laboratory for Conservation and Utilization of Subtropical Agro-bioresources, College of Life Science and Technology, Guangxi University, Nanning 530004, China; College of Horticulture and Landscape Architecture, Southwest University, 400715, Chongqing, China

**Keywords:** Allotriploid, subgenome asymmetric evolution, homologous exchange, Foc-TR4, carotenoids, starch

## Abstract

Bananas (*Musa* spp.) are one of the most important tropical fruits and staple food, which are of great significance to human societies. Plantain and Silk are two important banana subgroups, which are both triploid hybrids (AAB) between the wild diploid *Musa acuminata* and *M. balbisiana*. In this study, we reported the first haplotype-resolved genome assembly of Plantain and Silk bananas with genome size of approximately 1.4 Gb. We discovered widespread asymmetric evolution in the subgenomes of Plantain and Silk, which could be linked to frequent homologous exchanges (HEs) events. This is the first study to uncover the genetic makeup of triploid banana and verify that subgenome B harbors a rich source of resistance genes. Of the 88,078 and 94,988 annotated genes in Plantain and Silk, only 58.5% and 59.4% were present in all three subgenomes, with >50% genes containing differently expressed alleles in different haplotypes. We also found that Plantain is more resistant to banana Fusarium wilt, exhibiting a much faster defense response after pathogenic fungi infection. Many differentially expressed genes in abscisic acid, ethylene, jasmonic acid and salicylic acid pathways were identified in Plantain. Our analysis revealed that MpMYB36 promotes the biosynthesis of secondary cell wall and deposition of lignin by directly binding to the promoter of MpPAL and MpHCT, which allows Plantain to inhibit the penetration of early infection. Moreover, the insertion of the key carotenoid synthesis gene (*CRTISO*) may be the potential genetic basis for the richness of carotenoids in Plantain. Our study provides an unprecedented genomic basis for basic research and the development of elite germplasm in cultivated bananas.

## INTRODUCTION

Bananas (*Musa* spp.) are the largest herbaceous plants, mainly grown in tropical and subtropical regions and are of great significance to human societies (Kema and Drenth, 2020). Among bananas, dessert varieties (such as Cavendish) are one of the most widely traded fruits globally (FAOSTAT 2021), while starchy cooking types (like Plantains) are essential staples that contribute significantly to the diets of many developing countries (Robinson and Sauco, 2010). Most cultivated bananas are seedless triploid varieties (2n = 3× = 33) that were created through intra or inter-specific hybridization of the two Musa species, *M. acuminata* (A genome) and *M. balbisiana* (B genome) (Simmonds and Shepherd, 1955). Plantain, a crucial subgroup of cooking bananas, is a major dietary component for numerous populations in Africa, Latin America, and the Caribbean (Robinson and Sauco, 2010). In the major producing countries, per capita consumption of Plantain ranges from 40 kg/year in the Democratic Republic of Congo to 153 kg/year in Gabon (Akyeampong and Escalant, 1998). Genomic in situ hybridization (GISH) studies have confirmed that Plantain with an AAB genome has 21 A and 12 B chromosomes (D’Hont et al. 2000). Silk, a banana subgroup widely distributed in South and Southeast Asia, South America, and Australia, is a moderately vigorous plant that produces exceptionally flavorful dessert fruits with white flesh and a sub-acid, apple-like flavor. However, this subgroup is highly susceptible to Fusarium wilt, a devastating disease caused by *Fusarium oxysporum* f. sp. *Cubense* (Foc) (Dita et al., 2021; Zhan et al., 2022).

The complexities of assembling polyploid genomes stem from a variety of duplication events, including whole genome duplication (WGD) and segmental duplications that were recurrently observed in plant evolution. These duplications often result in the merging of repetitive sequences into a single collapsed region during assembly, which can lead to erroneous linkages with multiple genomic regions. The first draft genome assembly of *M. acuminata* spp. *Malaccensis* (DH-Pahang) was published in 2012 (D’Hont et al., 2012), which was subsequently refined in 2016 (Martin et al., 2016), and ultimately culminated in a telomere-to-telomere assembly (Belser et al., 2021). The *M. balbisiana* (DH-PKW) genome was assembled in 2019 (Wang et al., 2019). However, no cultivated banana genome has been sequenced for allotriploid up to now.

The complex and elusive nature of allotriploid genome sequences proves to be a challenge in investigating underlying molecular mechanisms for special traits, as allelic sequence variations are difficult to exclude. In this analysis, we presented our study of the haplotype-resolved genomes of two allotriploid cultivated bananas, Plantain and Silk. Our findings reveal the highly complex origin of the A subgenome in cultivated bananas, and a comparison of subgenomes A1, A2, and B helps us investigate genome evolution, genetic diversity, and functional divergence of subgenomes. In our transcriptome and functional analyses, we demonstrated that Plantain possesses a much faster defense response than Silk after Foc Tropical Race 4 (Foc-TR4) infection. We also examined the molecular difference underlying carotenoid production and starch metabolism in the two genomes by analyzing genomic and transcriptomic data from different developmental and postharvest stages. We discovered that the insertion mutation of a *CRTISO* gene in Plantain and the variation of gene number in Silk may be closely related to banana quality. Our findings overcome the limitations of allotriploid genome assembly, and provide a solid basis for understanding the origin, domestication, and genetic features of cultivated bananas.

## RESULTS

### Haplotype assembly and annotation of two AAB bananas genomes Plantain and Silk

Karyotype analysis confirmed that Plantain and Silk banana are highly complex allotriploid (2n = 3x = 33) genomes (Fig. 1a, Supplementary Fig. 1 and Supplementary Table 1) (Simmonds and Shepherd 1955). The genome size of Plantain and Silk was estimated to be ∼1.69 Gb and ∼1.52 Gb with a heterozygosity of 2.58% and 2.90%, respectively (Extended Data Fig. 1a). Plantain and Silk were sequenced separately using 59 Gb (35×) and 38 Gb (25×) PacBio HiFi reads, 272 Gb (161×) and 189 Gb (124×) PacBio CLR long reads, 167 Gb (98×) and 233 Gb (153×) Illumina reads and 205 Gb (121×) and 230 Gb (151×) Hi-C reads (Supplementary Table 2). Using the haplotype phasing and genome assembly pipeline presented in Figure 1b, we generated three haplotypes with contig N50 of 2.01∼2.92 Mb for Plantain and Silk, respectively (Cheng et al., 2022) (Table 1, Extended Data Fig. 1b and Supplementary Table 2). Moreover, over 90% of the Plantain reads and 93% of the Silk reads were anchored to final chromosomes (Supplementary Fig. 2 and Supplementary Table 3). The centromeric regions were 0.3∼3.7 Mb for Plantain and 0.3 ∼6.5 Mb for Silk, both of them contained 424 protein-coding genes (Supplementary Fig. 3 and 4, and Supplementary Table 4). Furthermore, more than half of the telomeres were identified in Plantain and Silk banana genomes (Table 1 and Supplementary Table 5).

**Figure 1.**
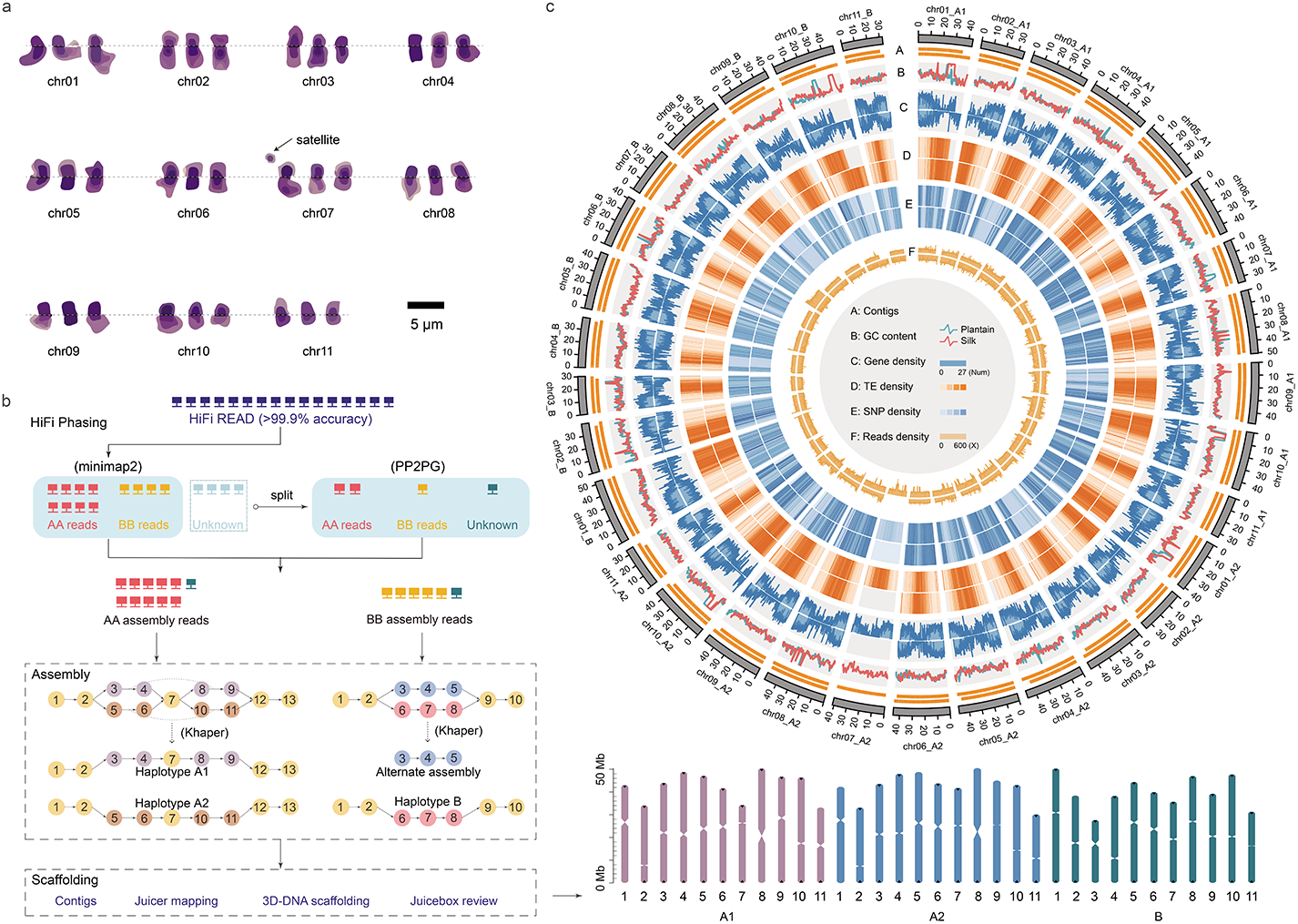
Overview of the Plantain (Batard) and Silk (Figue Pomme Géante) genomes assembly and features. **a,** Karyotype of Plantain with a scale of 5 μm. **b,** The flowchart of genome assembly and haplotype phasing. **c,** The circos diagram of Plantain and Silk. The circles from outer to inner separately represented contigs and gaps (A), GC (guanine cytosine) content (window size of 500 kb) (B), gene density (window size of 100 kb) (C), transposable element (TE) density (window size of 100 kb) (D), SNPs density (window size of 100 kb) (E), HiFi, CLR and Illumina reads coverage (window size of 100 kb) (F). For each track, the outer and inner layers indicate Plantain and Silk data, respectively.

The Long terminal repeat Assembly Index (LAI) (Ou et al., 2018) for Plantain and Silk were 14.69 and 19.69 respectively, with an average of ∼92% BUSCOs (Simão et al., 2015) Plantae reference genes in each assembly (Extended Data Fig. 1c and Supplementary Table 6). Consensus quality values (QV) were estimated using Merqury (Rhie et al., 2020), and were found to be 45.75 (96.65%) for Plantain and 42.16 (97.37%) for Silk (Table 1). Furthermore, the accuracy and completeness of the assemblies were supported by high mapping rates with PacBio HiFi reads, PacBio long reads, and Illumina reads (Supplementary Table 6). The subgenomes A and B of Plantain and Silk were highly consistent with the published *M. acuminata* (A genome) and *M. balbisiana* (B genome) genomes (Belser et al., 2021; Wang et al., 2019), and the collinearity among subgenomes was highly consistent (Extended Data Fig. 1de and Supplementary Fig. 5). The phasing accuracy of haplotype A1 and A2 in Plantain and Silk was confirmed by PCR results of alleles (Extended Data Fig. 1f and Supplementary Table 7). All of these results suggest that the assemblies for Plantain and Silk are of high quality.

The two assembled genomes of Plantain and Silk contained 56.47% and 54.37% transposable elements (TEs), respectively, which is consistent with other banana varieties of the *Musa* genus (Supplementary Table 8). TEs were predominantly found in intergenic regions, accounting for 74.90% and 72.35%, respectively, while only 3.69% and 3.57% were found in exonic regions in Plantain and Silk. Compared to diploid genomes, the insertion time of intact-LTRs in AAB genomes was later and the number was greater, suggesting that transposons in triploid bananas are more active (Supplementary Fig. 6). The Plantain and Silk genomes had 12,885 and 12,069 intact LTR-RTs, respectively, of which 66.74% and 66.49% had insertion times between 0 and 1 million years. This time frame is later than the divergence of *Musa acuminata* and *M. balbisiana* genotypes, which may have driven recent gene duplication and banana domestication (Supplementary Fig. 6 and Supplementary Table 9).

Plantain and Silk contained 88,078 and 94,988 protein-coding genes, respectively, with an average coding sequence length of approximately 1.2 kb and an average of five exons per gene (Supplementary Fig. 7 and 8 and Supplementary Table 10). Functional information was available for 97.84% and 98.28% of the genes in Plantain and Silk, respectively. In addition, 30,346 and 31,267 non-coding RNAs were annotated in Plantain and Silk, respectively (Supplementary Fig. 9 and Supplementary Table 11). The identified nucleotide-binging domain-like receptors (NLRs) in each accession consisted mostly of CNLs, followed by NLs, RNLs, and TNLs, with an uneven distribution across the chromosomes (Supplementary Fig. 10abc and Supplementary Table 12). Furthermore, a total of 435 and 481 putative WRKY genes were identified in Plantain and Silk (Supplementary Table 12), with high expression levels observed in rhizome, root tips, and root, particularly in response to Foc-TR4 infection (Supplementary Fig. 10d).

### Phylogenetic relationships of Musaceae and the ancestors of Plantain and Silk bananas

We constructed a phylogenetic tree of the Musaceae to clarify the evolutionary position of Plantain and Silk in the family (Fig. 2a and Supplementary Table 13). Our findings indicated that Pa1/Pa2 is more closely related to *M. acuminata* spp. *Banksia* compared to Sa1/Sa2, possibly due to variety differences. The subgenomes B (Sb and Pb) and *M. balbisiana* were found to be in the same clade, which supported previous studies that Plantain and Silk originated from a cross between the AA and BB genomes (Cenci et al., 2021). Functional enrichment analysis revealed that expanded gene families in Plantain were enriched in ‘protein kinase activity’, ‘transferase activity’, and ‘response to stress’, while expanded gene families in Silk were enriched in ‘organic substance biosynthetic process’, ‘phosphorus metabolic process’ and ‘protein metabolic process’ (Supplementary Fig. 11a and Supplementary Table 14). Notably, expanded gene families in Plantain were closely related to stress resistance compared to Silk.

**Figure 2.**
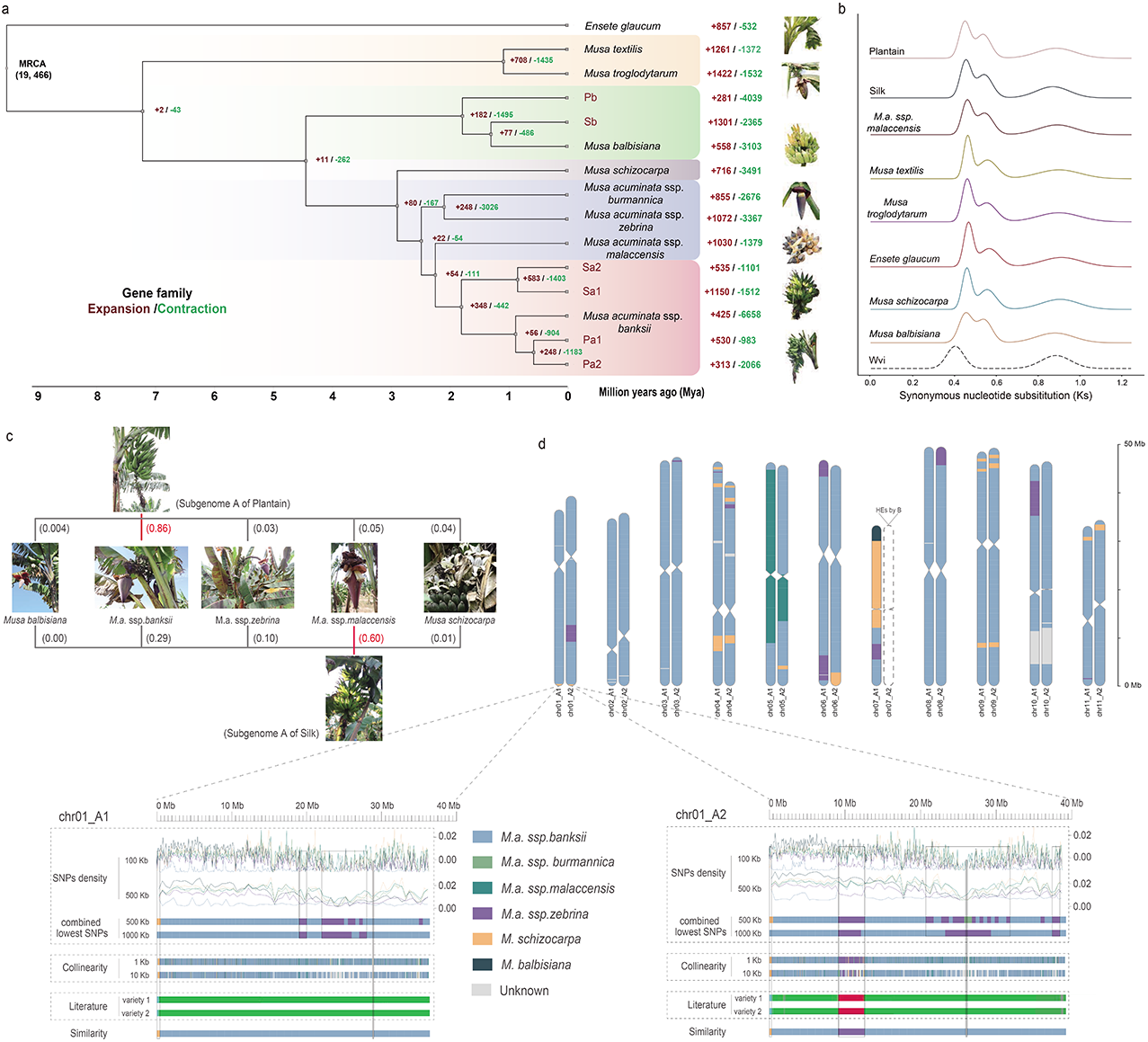
Phylogenetic relationships of Musaceae and genome ancestry mosaics for triploid cultivars Plantain and Silk. **a,** Phylogenetic tree of Plantain, Silk and other 9 Musaceae species (*Ensete glaucum* (Snow banana); *M. textilis* (Abaca); *M. troglodytarum* (Utafun); *M. balbisiana* (DH-PKW); *M. schizocarpa* (Schizocarpa); *M. acuminata* ssp*. burmannica* (Calcutta 4); *M. acuminata* ssp*. zebrina* (Maia oa); *M. acuminata* ssp. *malaccensis* (DH-Pahang); *M. acuminata* ssp. *banksii* (Banksii)) including their divergence time based on orthologues of the single gene family. **b,** Density distributions of the Ks values for homologous genes. Wvi represents *Wurfbainia villosa*. **c,** The pedigree composition of subgenome A in the triploid cultivated bananas Plantain and Silk. **d,** Chromosome ancestry painting of Plantain. The contributions of ancestral groups are represented along the 11 chromosomes by segments of different colors (green is *M. acuminata* ssp*. banksii*, red is *M. acuminata* ssp. *zebrina* and blue is *M. schizocarpa* of Literature).

Whole-genome duplications (WGDs) have played a significant role in angiosperm genome evolution. Previous studies suggest that Musaceae underwent three species-specific WGD events, namely the α/β and γ events (Lescot et al., 2008). After analyzing the Ks peak in pairwise genome comparison, we speculated that the α/β event occurred at about 58.67-59.67 Mya (Ks=0.528-0.537), which were different from the WGDs occurring in *W. villosa* from Zingiberaceae (Yang et al., 2021), and the γ event occurred at 98.56-100 Mya (Ks=0.887-0.900) shared in both Musaceae and Zingiberaceae (Fig. 2b, Supplementary Fig. 11b). As the occurrence time of the α/β WGD events was relatively close, the Ks of collinear block could not be entirely separated. However, we observed a collinear region on chromosomes 3, 6, 10, and 11 in subgenome A1 of Silk when Ks was ∼0.5, and most paralogous gene clusters shared relationships with three other clusters in all subgenomes, indicating that more than two WGDs had occurred (Supplementary Fig. 12).

Understanding patterns of interspecific introgression can reveal the origins of cultivated bananas (Martin et al., 2023). We precisely characterized the ancestral contributions of Plantain and Silk by examining the ancestry mosaics along the genome (Extended Data Fig. 2, Extended Data Fig. 3 and Supplementary Fig. 13). Both Plantain and Silk had at least five possible contributors of subgenomes A. For Plantain, we observed a dominant contribution (85.54%) from *M. acuminata* ssp. *banksii*, along with introgressions from *M. acuminata* ssp. *malaccensis* (5.07%), *M. acuminata* ssp. *zebrina* (3.11%), *M. schizocarpa* (4.06%) and *M. balbisiana* (0.36%). Silk, on the other hand, originated dominantly (59.86%) from *M. acuminata* ssp. *malaccensis*, with regions of *M. acuminata* ssp. *banksii* (29.22%), *M. acuminata* ssp. *zebrina* (9.55%), and *M. schizocarpa* (1.06%) (Fig. 2c and Supplementary Table 15). These results indicate that subgenome A underwent an extremely complex process of hybridization. Notably, we did not observe any *M. acuminata* ssp. *burmannica* contributions in Plantain and Silk triploids, and the subgenomes B of them were found to be homogenous (Fig. 2d and Supplementary Fig. 14). Our findings highlight that the origin of cultivated bananas is more complex than expected, involving multiple hybridization steps.

We further investigated genomic variations of the two AAB genomes. Overall, the genome sequence alignment between Plantain and Silk revealed high collinearity (Supplementary Fig. 15a). We found a total of 12,127,733 SNPs and 1,699,094 indels between Plantain and Silk, with an average of approximately 8.42 SNPs and 1.18 InDels per kilobase (Supplementary Table 16). The distributions of SNPs and InDels were positively correlated and both were more abundant in intergenic regions (Supplementary Fig. 15b). We identified 84.70 Mb as inversions between Plantain and Silk (Supplementary Table 17), and confirmed the authenticity of three inversions using PacBio HiFi reads to align to the assemblies (Supplementary Fig. 16). Between the haplotypes of Plantain and Silk, we found 105-255 and 142-240 inversions, and 55.81 Mb translocations were identified, with 7,435 inter-chromosomal translocations and 4,148 intra-chromosomal translocations (Supplementary Fig. 17 and Supplementary Table 17). We further characterized 3,886∼17,234 regions with cumulative lengths of 11.04∼67.14 Mb identified as PAVs, and these regions were associated with 743∼4,262 genes (Supplementary Fig. 18 and Supplementary Table 17, 18). KEGG enrichment analysis showed that ‘messenger RNA biogenesis’ and ‘starch and sucrose metabolism’ were mainly enriched (Supplementary Fig. 19). These findings may contribute to the quality of bananas.

### Asymmetric evolution between subgenomes in the allotriploid genomes

Loss of redundant genes is a common phenomenon that occurs following polyploidy (Zhao et al., 2017). We observed that gene loss regions overlapped significantly with homologous exchange (HE) regions, suggesting that loss of chromosomal segments after HEs is a key factor for gene loss (Fig. 3a, Extended Data Fig. 4 and Supplementary Fig. 20). Compared to *M. acuminata* ssp. *malaccensis* and *M. balbisiana*, Plantain lost 6,508 genes (3,463 in Pa and 3,045 in Pb), while Silk lost 5,237 genes (2,917 in Sa and 2,320 in Sb), more genes were lost in subgenome A than subgenome B (Supplementary Fig. 21a, and Supplementary Table 19). Specifically, the WRKY33 gene family, an important disease resistance gene family (Zhou et al., 2022), was lost more in subgenome A than B (Fig. 3b). Upon manual inspection of individual missing genes, we found that only 17.12% to 50.75% of the gene losses were complete absence, whereas 25.06% to 37.85% of the losses were altered genes caused by SNPs/InDels/TEs, and the remaining 24.19% to 45.02% gene losses were simply defined as ‘losses’ as they were not annotated as genes due to lack of expression (Supplementary Fig. 21b).

**Figure 3.**
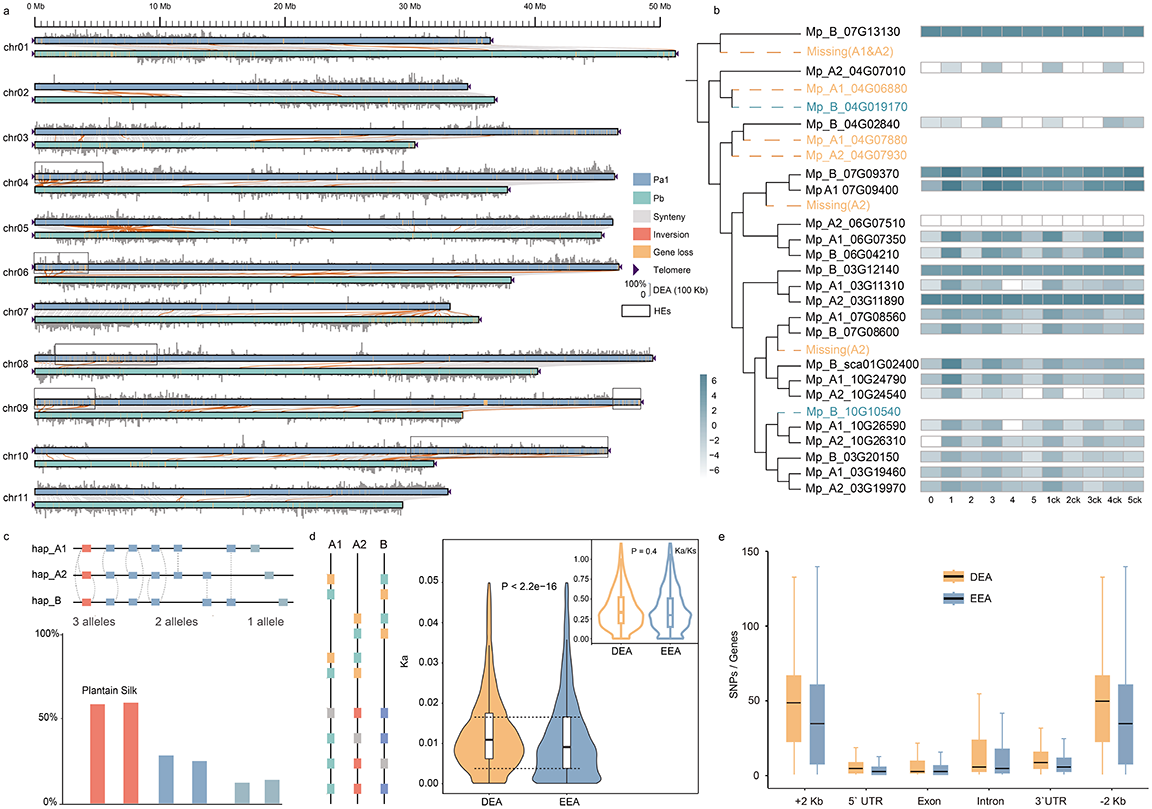
Subgenomic differentiation, asymmetric fractionation and expression of haplotypes. **a,** Asymmetry analysis between subgenome Pa1 and Pb. All gene loss regions between Pa1 and Pb are shown in yellow blocks. DEA percentage distribution is plotted above or under each chromosome in 100-kb bins. The area framed by the box are HEs. The black triangles indicate the presence of telomere sequence repeats. The collinear regions between Pa1 and Pb are shown linked by gray lines. **b,** The picture shows the evolutionary tree of the WRKY33 gene family in Plantain, and its changing hot map of the expression (log2 (TPM)) in each period after the pathogen is infected. In the evolution, Missing represents the loss of genes in A1 or A2 subunit, and the brown font represents genetic variations resulting in functional changes. **c,** Identification strategies and statistics for alleles. Allelic gene pairs were selected according to the following rules: (1) paired regions must be on homologous haplotypes; (2) when there is one-to-many paired genes, take the one with the higher C-score (score(A, B) / max(score(A,), score (,B))); (3) the three genes are paired with each other are identified as 3 alleles, the two genes are paired with each other are identified as 2 alleles, the others are 1 allele; (4) Syntenic gene pairs defined above were double-checked manually. **d,** DEAs has relatively higher Ka and Ka/Ks value than EEAs in Plantain. P-values were calculated with two-sided Student’s t-test. **e,** SNPs density in DEAs/EEAs features of Plantain.

Based on the phased haplotypes, 58.46% and 59.37% of annotated genes were present in all three subgenomes, while 28.25% and 25.01% in were present two subgenomes, and 12.18% and 13.91% were present in one subgenome, with an average of 2.44 and 2.42 copies per gene in Plantain and Silk, respectively (Fig. 3c and Supplementary Table 20). To assess the rate of evolution on alleles, we calculated Ka and Ks values between allelic pair, and the vast majority of alleles Ka/Ks were low (<0.05) (Supplementary Fig. 22). About 3.81% and 4.35% (3,352 and 4,132) of allelic pairs showed possible positive selection (Ka/Ks>1) (Supplementary Table 21). Consistent with previous analyses on the effects of copy number variations (CNVs) on gene expression (Pham et al., 2017), we observed a positive correlation between allelic copies and gene expression (Supplementary Fig. 23). To investigate the homologous gene expression patterns and their divergence in the three subgenomes, we compared the genome-wide transcriptional levels of subgenomes A and B based on 17,162 and 18,799 homologous gene pairs in different tissues of Plantain and Silk (Supplementary Fig. 24). A total of 9,014 and 10,015 homoeologous gene pairs (∼62.73% and ∼64.04%) had expression difference larger than 2-fold change in at least one tissue, including 4,669/5,774 and 5,430/5,578 homoeologs having higher expression in subgenomes A and B, respectively. Among these homoeologs, 3,584/4,437 and 4,345/4,241 had higher expression values exclusively in all tissues of subgenomes A and B, respectively, while 1,085 and 1,337 homologs had swinging expression bias (Supplementary Table 22). The homologous expression bias showed asymmetric expression patterns between subgenomes A and B in Plantain and Silk.

After investigating allelic imbalance, which refers to the differential expression of alleles (DEAs), we observed a log-linear increase in DEAs with the number of RNA-seq samples, leveling off at over 35 and 23 samples of Plantain and Silk, respectively (Supplementary Fig. 25ab and Supplementary Table 23). A total of 52,338 and 49,388 DEAs were identified in Plantain and Silk, respectively (Fig. 3d, Supplementary Fig. 25c and Supplementary Table 24), with 25.76% and 23.97% showing significant expression differences among three alleles (Supplementary Fig. 26). Notably, DEAs exhibited significantly higher Ka (t-test, P value=2.2×10^−16^) and Ka/Ks (t-test, P value=2.0×10^−7^) than equivalently expressed alleles (EEAs), indicating a potentially faster evolutionary rate of DEAs (Fig. 3d). Additionally, the promoter, exon, intron, 5’ UTR, and 3’ UTR regions of DEAs had higher SNP densities than EEAs, which may lead to the difference of their expression (Fig. 3e and Supplementary Fig. 25d).

We also investigated the expression of nucleotide-bounding leucine-rich repeat proteins (NLRs), WRKY22 and leucine-rich repeat receptor-like kinase (LRR-RLK) resistance gene families, and genes involved in carotenoid synthesis and the ethylene pathway in different subgenomes. Notably, the expression of NLRs, WRKY22, and LRR-RLK resistance genes was higher in subgenome B than in subgenome A, which may be attributed to variations between alleles (Extended Data Fig. 5a-f and Supplementary Fig. 27). Furthermore, the expression of carotenoid pathway genes was higher in subgenome A, particularly in Silk’s subgenome A, where it was three times higher than in B during the decomposition period, potentially leading to the rapid reduction of Silk’s carotenoid accumulation. In contrast, during the ripening period, the expression of ethylene pathway genes was higher in subgenome A than in B, and higher in Plantain than in Silk (Extended Data Fig. 5g-n). These findings suggest that asymmetric evolution has significantly impacted the genetic basis of banana resistance, with subgenome B contributing more and subgenome A being more involved in carotenoid degradation and ethylene ripening. This distinction is also reflected in the phenotypes of ancestral species (Wang et al., 2019), providing valuable resources and guidance for genome-based molecular marker-assisted breeding of bananas.

### Plantain has a faster response to Foc-TR4 than Silk by comparing transcriptome

We conducted field and pot experiments to assess the difference in resistance to Foc-TR4 infection between Plantain and Silk. Our observations indicated that Silk exhibited typical infection symptoms, such as leaf yellowing and pseudostem splitting, while Plantain showed no discernible signs of infection (Fig. 4a and Supplementary Fig. 1). The average Rhizome Discoloration Index (RDI) for Plantain was 1, whereas Silk had an RDI of 3.7 (Fig. 4b, Supplementary Fig. 28 and Supplementary Table 25). These findings confirm that Plantain displays high resistance to Banana *Fusarium* wilt, while Silk is highly susceptible to it.

**Figure 4.**
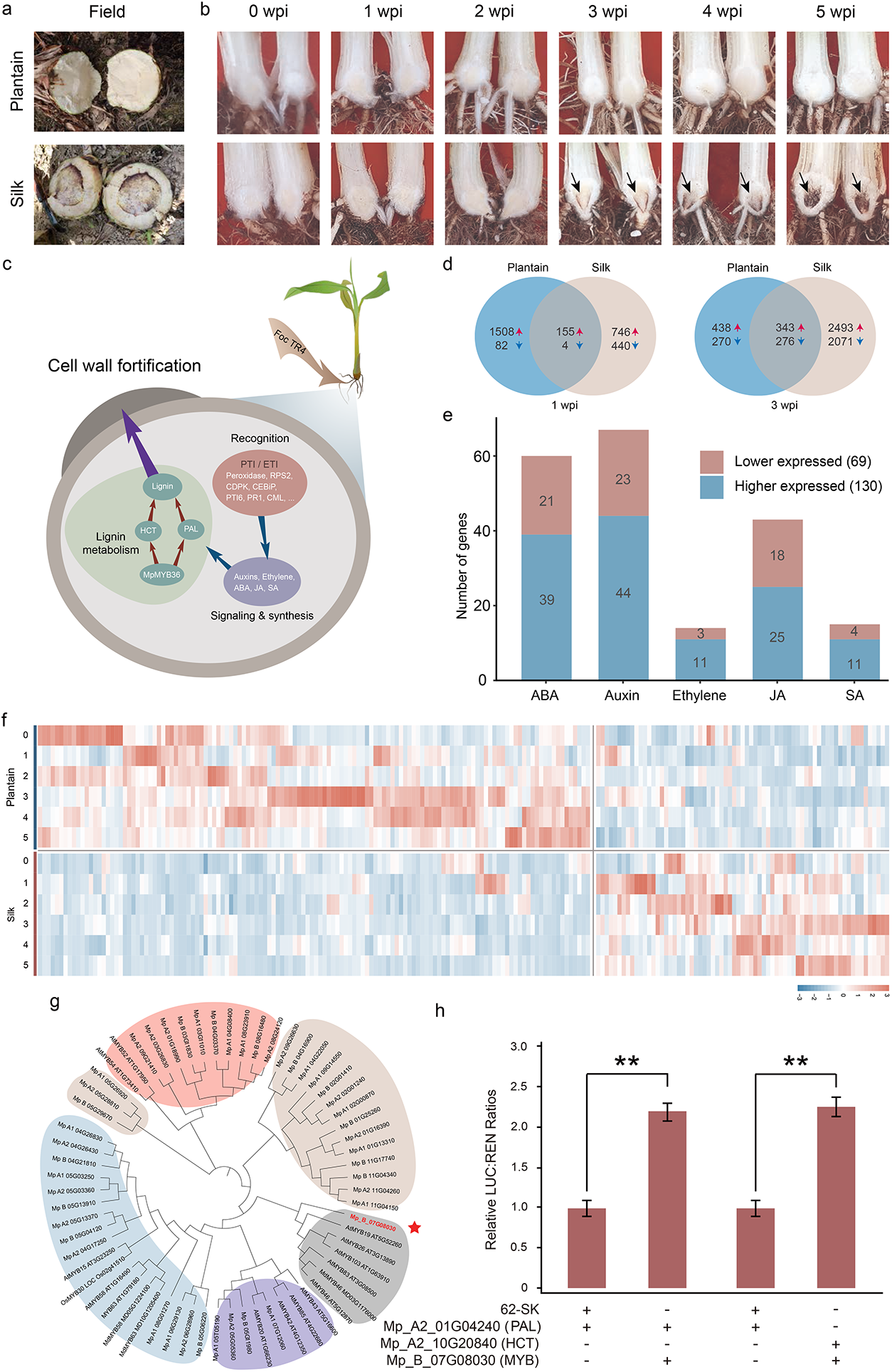
Resistance and mechanism differences of *Fusarium oxysporum* f. sp. *cubense* Tropical Race 4 between Plantain (Batard) and Silk (Figue Pomme Géante). **a,** The rhizome of the Plantain and Silk inoculated with Foc-TR4 in the field. A deep golden discoloration of the inner rhizome develops of Silk. **b,** The rhizome of Plantain and Silk, which cut in half longitudinally, inoculated with Foc-TR4. Plantain’s rhizome showed no traces of brown discoloration in the lower and center regions, while Silk’s rhizome developed extensive brown discoloration from 3 wpi, which associated with high Foc sensitivity. **c,** Schematic representation of the response of banana against Foc-TR4. **d,** Venn diagrams of differentially expressed genes at 1 wpi and 3 wpi of Plantain and Silk. **e,** Pathway distribution of the 199 differentially expressed genes involved in plant hormone signal transduction. Blue color indicates higher and orange color indicates lower relative expression in Plantain compared with Silk. **f,** Heatmap of genes at 1-5 wpi after inoculated with Foc-TR4 which from e. **g,** Phylogenetic tree of differentially expressed MYBs in Plantain with other known cell wall-associated MYB transcription factors. **h,** Mp_B_07G08030 (MYB) enhanced fluorescence intensity of LUC driven by the Mp_A2_01G04240 (PAL) and Mp_A2_10G20840 (HCT) promoter compared to the control. The mean ± s.d. of three biological replicates is shown.

To gain a better understanding of the mechanism behind the resistance difference between Plantain and Silk, we identified differentially expressed genes (DEGs) at 0, 1, 2, 3, 4 and 5 weeks post-inoculation (wpi) following Foc-TR4 infection (Supplementary Table 26). At 1 wpi of Foc-TR4, the number of up-regulated genes identified in Plantain (1,663) was significantly higher than that in Silk (901). However, from 3 wpi onwards, the number of DEGs identified in Silk increased rapidly, and there were few shared DEGs between Plantain and Silk at 4 and 5 wpi (Fig. 4d and Supplementary Fig. 29). KEGG enrichment analysis showed that at 1 wpi, the DEGs of Plantain were highly enriched in well-known resistance pathways, such as “plant-pathogen interaction”, “plant hormone signal transduction”, and “phenylpropanoid biosynthesis” (Supplementary Fig. 30). In contrast, Silk’s DEGs were enriched in some metabolic pathways unrelated to disease resistance at 1 wpi, but this trend was reversed by 3 wpi (Supplementary Table 27). Overall, Plantain exhibited a significantly faster response to Foc-TR4 infection than Silk.

To investigate the genes involved in plant-pathogen interactions, including pathogen-associated molecular pattern (PAMP)-triggered immunity (PTI) and effector-triggered immunity (ETI), which constitute the first layer of plant defense response that restricts pathogen proliferation, we identified candidate DEGs in Plantain and Silk (Supplementary Fig. 31). These DEGs included peroxidase, RPS2, CDPK, CEBiP, PTI6, PR1, and CML, of which seven were verified by RT-qPCR, showing consistent trends with the RNA-seq analysis (Supplementary Table 28). We also examined genes involved in phytohormone signaling and response, identifying 199 DEGs across six time points, representing various plant hormone signaling and response pathways, such as auxin, abscisic acid (ABA), ethylene, jasmonic acid (JA), and salicylic acid (SA) (Supplementary Table 29). Among them, 130 genes (65.33%) were expressed higher in Plantain than Silk (Fig. 4e). These findings suggest that banana response to Foc-TR4 infection involves multiple phytohormone signaling pathways and responses (Fig. 4f).

Phenylpropionic acid biosynthesis and flavonoid biosynthesis are part of the secondary metabolism and play an important role in plant defense by strengthening cell walls and producing phytoalexins. We identified the expression of lignin pathway genes in Plantain and Silk, and found that the responses of PAL, 4CL, HCT, CCoAOMT, CCR, CAD, and POD/LAC genes in Plantain to Foc-TR4 infection were much faster than those in Silk (Supplementary Table 30). Furthermore, we observed that Plantain had a higher number of F5H and COMT genes compared to Silk (Extended Data Fig. 6). The chalcone synthase (CHS) gene (Ferrer et al., 1999), which is crucial for the biosynthesis of flavonoid antibacterial phytocyanins and anthocyanin pigments in plants, was expressed earlier in Plantain and had more copies than in Silk (Extended Data Fig. 6). These findings suggest that the expression of genes involved in the phenylpropane biosynthesis pathway was activated earlier in Plantain than in Silk in response to Foc-TR4 infection. Additionally, we found that the DEGs of several common resistance gene families in Plantain were mostly identified at 1 wpi after Foc-TR4, while in Silk, they did not appear until 3 wpi (Supplementary Fig. 32 and Supplementary Table 31).

MYB genes encodes a large family of transcription factors (TFs) that play an important role in the regulation of lignin synthesis (Dubos et al., 2010). Phylogenetic analysis revealed that MpMYB36 (Mp_B_07G08030) belonged to the same subfamily as AtMYB46, a second-layer master switch of secondary cell wall biosynthesis (Zhong et al., 2007). Most of the MYBs in this cluster (Fig. 4g) have been shown to be involved in lignin biosynthesis (Zhong et al., 2007; Chen et al., 2019; McCarthy et al., 2009; Yang et al., 2007). Co-expression network analysis indicated that 34 differentially expressed MYBs in Plantain were grouped with lignin biosynthesis genes into distinct co-expression clusters after Foc-TR4 infection (Supplementary Fig. 33a and Supplementary Table 32). Among these MYBs, MpMYB36 was positively correlated with 33 lignin biosynthesis genes (Supplementary Fig. 33b). To clarify the role of MpMYB36 in promoting deposition of lignin in the secondary cell walls, we analyzed promoter regions of 11 lignin biosynthesis-related genes. At least one AC/secondary wall MYB-responsive element was identified in the promoter region of each gene (Supplementary Fig. 34). Next, we performed a transient expression assay using a dual-luciferase reporter system, which showed that the co-expression of MpMYB36 (non-empty vector control) with LUC driven by the Mp_A2_01G04240 (PAL) and Mp_A2_10G20840 (HCT) promoter significantly increased the LUC/REN ratio (Fig. 4h), demonstrating that MpMYB36 directly up-regulated the expression of Mp_A2_01G04240 and Mp_A2_10G20840. Based on these results, we created a simple schematic illustration of the plant defense against Foc-TR4 infection in banana (Fig. 4c).

### Genomic insights into carotenoid synthesis and starch metabolic in cultivated banana

Carotenoids are abundant in bananas, some of which can be converted into vitamin A in human body. To study the difference of carotenoids content between Plantain and Silk, we collected RNA samples from five fruit developmental stages (Fig. 5a and Supplementary Fig. 35), and found that the carotenoid content of Plantain was higher than that of Silk during the same growth period (Extended Data Fig. 7a). A total of 44 and 48 key genes involved in the metabolic synthesis pathway of carotenoids were identified in Plantain and Silk, respectively (Supplementary Table 33). The expression of ’early’ biosynthesis genes was greater than that of ’late’ biosynthesis genes in the carotenoid synthesis pathway, and the synthesis genes of Plantain had higher expression levels than those of Silk in the ’early’ biosynthesis stage (Fig. 5a, Extended Data Fig. 7b and Supplementary Table 34), suggesting that the early synthetic pathway had a greater impact on carotenoid synthesis. We also found that *CRTISO* genes were predominantly expressed in Plantain, and carotenoids were more accumulated in Plantain than Silk (Fig. 5b, Supplementary Fig. 36 and 37). For *CRTISOs* of Plantain, the coding regions of two genes in subgenome A (*CRTISO1* and *CRTISO2*) were identical compared with *CRTISO3* in subgenome B, there was an 87 bp insertion (between the 5th and 6th exon) in these two *CRTISO* genes of subgenome A, and this insertion was absent in *CRTISO* genes of Silk, *Musa acuminata* (Belser et al., 2021), and *Musa balbisiana* genomes (Fig. 5c, Extended Data Fig. 7c and Supplementary Fig. 38) (Wang et al., 2019). In addition, we found that the binding amino acids of lycopene of *CRTISO1* gene in Plantain and Silk have changed. At the same time, during the kinetic simulation of the binding of *CRTISO1* and lycopene, the stability of the binding system is also quite different (Supplementary Fig. 39). *CRTISO1* and *CRTISO2* with this insertion were also expressed at higher levels than the homologous genes in Silk, suggesting that this insertion may make the *CRTISO* gene in Plantain more active, thereby affecting the carotenoid synthesis pathway.

**Figure 5.**
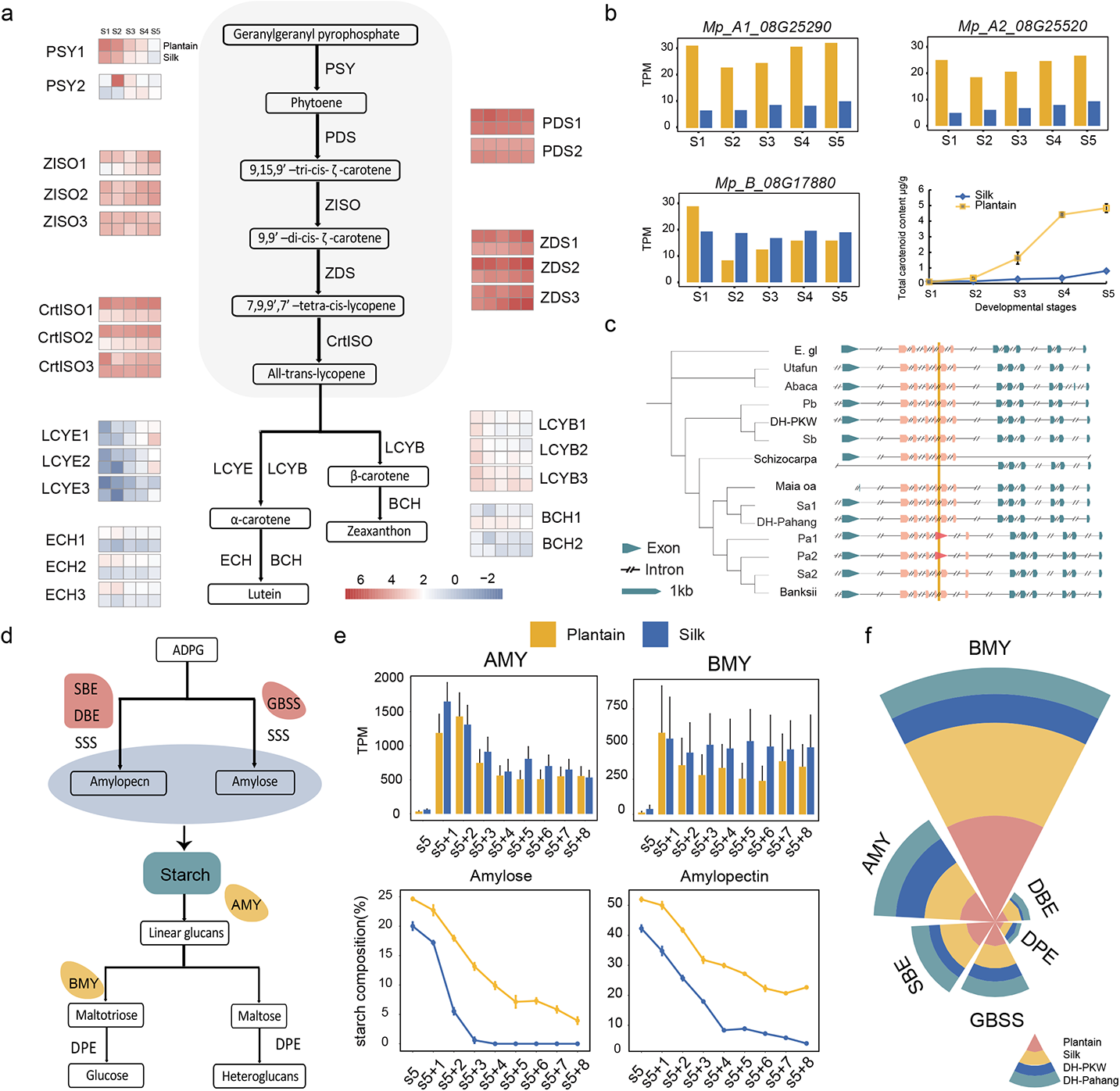
Multi-omics differential analysis of carotenoids and starches metabolism. **a,** Overview of the carotenoid synthesis pathway. Genes aligned horizontally in the heatmap indicate genes at five developmental stages in Plantain genomes. We divided the carotenoid synthesis pathway encoding genes into two groups designated as ‘early’ (grey background) and ‘late’, respectively. Low to high expression is indicated by a change in color from blue to red. PSY: phytoene synthase; PDS: phytoenedesaturase; ZISO: ε-carotene isomerase; ZDS: ε-carotenedesaturase; CRTISO: carotenoid isomerase; LCYE: lycopene δ-cyclase; LCYB: lycopene β-cyclase; BCH: β-carotene hydroxylase; ECH: ε-carotene hydroxylase. **b,** The barplot graph shows the gene expression profile (TPM) of CRTISO1 at five developmental stages in Plantain and Silk. The line graph shows changes of all carotenoid content. **c,** CDS sequence comparison of CRTISO in 11 species (Egl: *E. glaucum*; SY137: *M. troglodytarum*; U9: *Musa* textilis; BB: *M. balbisiana*; SS: *M. schizocarpa*; Bur: *M. acuminata* spp. *burmannica*; Zeb: *M. acuminata* spp. *zebrina*; AA: *M. acuminata* spp. *malaccensis*; Ban: *M. acuminata* spp. *banksii*). **d,** Overview of the starch synthesis and degradation pathway. GBSS: Granule-bound starch synthase; SSS: Soluble starch synthase; SBE: Starch branching enzyme; DBE: Starch debranching enzyme. AMY: α-amylase; BMY: β-amylase; DPE: Starch phosphorylase. **e,** The bar graph shows the gene expression profile (TPM) at eight postharvest stages in Plantain and Silk. The other two graphs below show the starch hydrolysis of the fruit at 8 stages after harvest. **f,** The circular bar gragh shows the gene numbers of 6 types of genes of 4 species (Plantain, Silk, AA and BB) related to starch metabolism.

Bananas are a high-starch fruit with a high ratio of amylose to amylopectin, and they can synthesize resistant starch after heat moisture treatment, thereby improving the structure of the gut microbiota in the human body (Villa, 2020). To study the difference in starch content between Plantain and Silk, we collected bananas at five developmental stages (S1-S5) and eight postharvest stages for comparative analysis. We identified 90 starch metabolism-related genes in the Plantain genome, including 28 in the starch synthesis pathway and 62 in the starch degradation pathway (Fig. 5d). Similarly, 98 such genes were identified in the Silk genome, including 30 in the starch synthesis pathway and 68 in the starch degradation pathway (Supplementary Table 35). During the early stage of starch synthesis, the average expression levels of genes related to starch synthesis in Plantain were higher than those in Silk. We found that starch accumulation mainly occurred in the early stage (S1-S3), with the peak in S4-S5 stage (Supplementary Figure 40). Regarding the starch degradation pathway, the number of β-amylase genes in Plantain and Silk was significantly higher than that in *M. acuminata* and *M. balbisiana* (Fig. 5f; Supplementary Table 36). Furthermore, the mean expression levels and gene number of BMY in Silk were higher than in Plantain (Fig. 5e). Notably, the degradation rate of both amylopectin and amylose was faster in Silk than in Plantain (Fig. 5e, Supplementary Table 37 and 38). Therefore, genomic and transcriptome analysis revealed that the number of genes related to starch degradation in Silk and their gene expression levels were higher than those in Plantain.

## DISCUSSION

Previously, most genomes were mosaic assemblies, however, this approach results in significant loss of information for highly heterozygous polyploid species. Developing a haplotype-resolved genome for such species remains a challenge. Currently, there are four strategies for phasing: 1) The initial contig assembly followed by identification and duplication of collapsed contigs based on read depth, the augmented set of sequences was subjected to haplotype phasing along with initial phased contigs, resulting in a fully haplotype-solved assembly (Zhang et al., 2021); 2) Trio binning (Koren et al., 2018), that can recover both parental haplotypes from F1 individuals by partitioning parental unique reads before assembly, but this method is time-consuming and laborious, which is not conducive to popularization; 3) Inferring regional haplotypes by aligning sequenced reads to a reference genomes (Chin et al., 2016), however, these efforts are limited by the continuity of an available reference assembly; 4) High-throughput/resolution chromosome conformation capture (Hi-C) technology has helped to provide allele-resolved assemblies (Zhang et al., 2019). With the help of ultra-high accuracy PacBio HiFi reads, CLR reads, Hi-C reads, Illumina short reads, the telomere-to-telomere gapless chromosomes of its ancestral species, and used a combination of assembly strategies 3 and 4, we firstly reported two haplotype-resolved assemblies of allotriploid cultivated bananas Plantain and Silk. The contig N50, GC content, full-length transcripts and other indices showed a high level of integrity and accuracy of the reference genomes. The first two haplotype-resolved genomes of AAB allotriploid bananas provided a basis for further genetic studies of *Musa*.

Our genome mosaics results demonstrated complex specific hybridization origins for Plantain and Silk, involving at least six ancestors. We found that their subgenomes A were more complex than expected, with Plantain being a Banksii-rich cultivar and Silk being a DH-Pahang-rich cultivar. The regions with unknown contributions indicate the existence of other unknown ancestors. Our results obtained by direct comparison among genomes are more accurate than those obtained by Illumina reads, the most intuitive show is that each locus is its true chromosomal location of Plantain and Silk. Recombination between A and B genomes was visible, confirming that several interspecific hybridization steps occurred at their origin, as previously suggested (Cenci et al., 2021). We also showed that the contribution of *M. schizocarpa*, previously thought to be restricted to a few *M. schizocarpa* × *M. acuminate* cultivars and suspected to be present in east African highland bananas, was present in Plantain and Silk. It is worth noting that there is a significant difference between the Silk reported in (Martin et al., 2023) and ours, so we speculate that they may not be the same Silk cultivar.

Musaceae species shared three WGD events, and the variation and loss of genome fragments resulting from whole genome duplication led to drastic changes in gene families across different species. Our results indicated that functional divergence of subgenomes occurred in polyploidy bananas after WGD. It is worth noting that homoeologous exchanges may obscure the signal of expression dominance in subgenomes of allopolyploids, which can result in a series of rapid genetic and epigenetic modifications for agronomic traits (Bird et al., 2018). Asymmetric subgenomic fractionation occurred in the allopolyploid, primarily by accumulation of small deletions in gene clusters through illegitimate recombination. We observed that gene loss regions highly overlap with HE regions, indicating that loss of chromosomal segments after HEs is one of the key factors in gene loss (Fig. 3a). Differentially expressed alleles (DEAs), which have profound effects on growth and evolvability. This could be due to the different distribution of SNPs at the promoter regions of adjacent genes that is associated with levels of gene expression. Asymmetric evolution significantly impacted the genetic basis of banana resistance, with subgenome B providing greater contributions and subgenome A being more involved in carotenoid degradation and ethylene ripening. These finding provide new resources and guidance for genome-based molecular marker assistant breeding for bananas.

*Fusarium* wilt, caused by Foc, is a destructive soil-borne fungal disease that severely threatens the sustainable development of global banana industry. While Foc-TR4 can cause severe yield losses in Silk, cooking bananas such as Plantain, appear to be resistant (Zuo et al. 2018). Our results confirmed that Plantain has a faster response than Silk (Fig. 4). KEGG enrichment analysis showed that DEGs in the first week after Foc-TR4 infection were highly enriched in well-known resistance pathways in Plantain including “plant-pathogen interaction”, “plant hormone signal transduction” and “phenylpropanoid biosynthesis” (Supplementary Fig 36). After Foc-TR4 infection, the expression of PTI and ETI genes increased in Plantain (Supplementary Fig 37). Many plant hormone signaling and response pathways, including auxin, ABA, ethylene, JA and SA, were higher expressed in Plantain than in Silk (Fig. 4e). These findings suggest that banana response to Foc-TR4 infection involves multiple phytohormone signaling pathways and responses (Fig. 4f). MYB transcription factors play an important role in the regulation of lignin synthesis (Dubos et al., 2010). We identified a MYB transcription factor located on chr07B of Plantain, which was highly positively correlated with 33 lignin biosynthesis genes (Supplementary Fig. 40). Double luciferase assay confirmed that the MpMYB36 (Mp_B_07G08030) directly regulated PAL (Mp_A2_01G04240) and HCT gene (Mp_A2_10G20840), thereby positively regulating the lignin pathway and participating in the response to TR4 infection (Fig. 4h).

Bananas are abundant in ascorbic acid (vitamins C), β-carotene (provitamin A), magnesium (Mg), and potassium (K) (Wall and Marisa, 2006). Carotenoids, present in chromoplasts, can endow flowers and fruits with their distinct coloration (Hirschberg, 2001). There are different greatly in carotenoid content of Plantain and Silk. The expression of carotenoid synthesis genes was much higher than decomposition genes in the developmental stage, indicating that carotenoid accumulation was crucial for fruit development. *CRTISO* is an important upstream synthetic gene in the carotenoid metabolic pathway, and there is large difference in gene structure between the two varieties, which may be related to the significantly higher expression level in Plantain than in Silk, thus affecting carotenoids of anabolic. Starch content varied in the range of 61.30–86.76% among different banana cultivars (Ravi and Mustaffa, 2013). Compared to Silk, Plantain contained more starch synthesis genes during the developmental period. After the fruit was picked and ripened, there were more amylolytic genes expressed in Silk than in Plantain, and the expression level was also higher than that in Plantain, which made the accumulation of starch in Silk significantly reduced, and greatly damaged its economic value. The content of carotenoids and starch in the fruits of Silk (Dessert) and Plantain (Cooking) varies widely, but the molecular regulation of the difference is unclear. So more detailed and in-depth research is needed to resolve this issue.

## METHODS

### Plant materials

Plantain and Silk were introduced from Centre Africain de Recherche sur Bananiers et Plantains (CARBAP) and International *Musa* germplasm Transit Center (ITC), respectively. Plantain is a French Horn type from the starchy plantain subgroup, which is one of the most popular cultivars in West Africa (De Langhe et al., 2005; Ibobondji et al., 2018). Silk (ITC0769, DOI: 10.18730/9KGW1), is a dessert cultivar bearing sweet acidic fruits with an apple-like flavor (Robinson and Sauco, 2010). Samples of the two cultivars were collected from National Center for Banana Genetic Improvement in Guangzhou, China.

### PacBio library construction and sequencing

The SMRT bell library target size to construct depends on the goals of the project and the quality and quantity of the starting gDNA. The g-TUBE can be used to shear gDNA fragments for constructing 10 -20 kb SMRT bell libraries. After shearing, AMPure PB Beads were used to concentrate sheared gDNA. ExoVII was used to treat DNA for shearing long overhangs before DNA damage repair. T4 DNA Polymerase was used to fill in 5’ overhangs and remove 3’ overhangs. And T4 PNK was used to phosphorylates 5’ hydroxyl group. Then, SMRT bell hairpin adapters included in Template Prep Kit are ligated to repaired ends. Next, we do size selection using Blue Pippin System and set size cut-off threshold depending on the goals of the project. Then, AMPure PB Beads were used to concentrate and purify SMRT bell templates after size selection. Sequencing primer annealed to both ends of the SMRT bell templates and polymerase is bound to both ends of SMRT bell templates using Binding Kit. Finally, use DNA Sequencing Reagent Kit and follow the manual to load libraries in SMRT Cells.

### Illumina short-reads sequencing

DNA degradation and contamination was monitored on 1% agarose gels. DNA purity was checked using the NanoPhotometer® spectrophotometer (IMPLEN, CA, USA). DNA concentration was measured using Qubit® DNA Assay Kit in Qubit® 2.0 Flurometer (Life Technologies, CA, USA). A total amount of 1.5µg DNA per sample was used as input material for the DNA sample preparations. Sequencing libraries were generated using Truseq Nano DNA HT Sample preparation Kit (Illumina USA) following manufacturer’s recommendations and index codes were added to attribute sequences to each sample. These libraries constructed above were sequenced by Illumina NovaSeq 6000 platform and 150 bp paired-end reads were generated with insert size around 350 bp.

### Hi-C library preparation and sequencing

Hi-C library construction following the standard protocol with certain modifications (Belton et al., 2012). After ground with liquid nitrogen and cross-linked by 4% formaldehyde solution at room temperature in a vacuum for 30 mins. 2.5 M glycine was added to quench the crosslinking reaction for 5 min and then put it on ice for 15 min. The sample was centrifuged at 2500 rpm at 4°C for 10 mins, and the pellet was washed with 500μl PBS and then centrifuged for 5 min at 2500 rpm. The pellet was re-suspended with 20μl of lysis buffer (1 M Tris-HCl, pH 8, 1 M NaCl, 10% CA-630, and 13 units protease inhibitor), the supernatant was then centrifuged at 5000 rpm at room temperature for 10 min. The pellet was washed twice in 100μl ice cold 1x NEB buffer and then centrifuged for 5 min at 5000 rpm. The nuclei were re-suspended by 100μl NEB buffer and solubilized with dilute SDS followed by incubation at 65°C for 10 min. After quenching the SDS by Triton X-100, an overnight digestion was applied to the samples with a 4-cutter restriction enzyme DPNII (GATC) (400 units) at 37°C on a rocking platform. The following steps involved marking the DNA ends with biotin-14-dCTP and blunt-end ligation of the cross-linked fragments. The proximal chromatin DNA was re-ligated by ligation enzyme. The nuclear complexes were revers cross-linked by incubation with proteinase K at 65°C. DNA was purified by the phenol-chloroform extraction. Biotin was removed from non-ligated fragment ends using T4 DNA polymerase. Ends of sheared fragments by sonication (200-600 bp) were repaired by the mixture of T4 DNA polymerase, T4 polynucleotide kinase and Klenow DNA polymerase. Biotin-labeled Hi-C samples were specifically enriched using streptavidin C1 magnetic beads. After adding A-tails to the fragment ends and following ligation by the Illumina paired-end (PE) sequencing adapters, Hi-C sequencing libraries were amplified by PCR (12-14 cycles) and sequenced on Illumina NovaSeq-6000 platform (PE 150 bp).

### RNA quantification and transcriptome sequencing

RNA degradation and contamination were monitored on 1% agarose gels. RNA purity was checked using the NanoPhotometer® spectrophotometer (IMPLEN, CA, USA). RNA integrity was assessed using the RNA Nano 6000 Assay Kit of the Bioanalyzer 2100 system (Agilent Technologies, CA, USA). A total amount of 1 µg RNA per sample was used as input material for the RNA sample preparations. Sequencing libraries were generated using NEBNext® UltraTM RNA Library Prep Kit for Illumina® (NEB, USA) following manufacturer’s recommendations and index codes were added to attribute sequences to each sample. The clustering of the index-coded samples was performed on a cBot Cluster Generation System using TruSeq PE Cluster Kit v3-cBot-HS (Illumia) according to the manufacturer’s instructions. After cluster generation, the library preparations were sequenced on an Illumina Novaseq platform and 150 bp paired-end reads were generated.

### Estimation of the genome size and heterozygosity

The genome size was estimated through k-mer frequency analysis, which involves analyzing the distribution of k-mers in the genome using Poisson’s distribution. Prior to assembly, we used Jellyfish (v2.2.7) (Marçais et al., 2011) to generate the 17-mer distribution of 167 Gb (Plantain) and 233 Gb (Silk) Illumina short reads, which we then uploaded to the GenomeScope website (http://qb.cshl.edu/genomescope/). This analysis revealed an estimated genome size of 1694.56 Mb with a 2.58% heterozygous rate for Plantain and 1520.59 Mb with a 2.90% heterozygous rate for Silk genome.

### Genome assembly

#### PacBio HiFi reads and Hi-C reads phasing

Based on published telomere-to-telomere AA and BB reference genomes, we phased PacBio HiFi reads and Hi-C reads of Plantain and Silk by aligning them to AA and BB reference genomes, which can be briefly summarized as follows: First, PacBio HiFi reads and Hi-C reads were mapped to the published AA and BB reference genomes through Minimap2 (2.18-r1015) (Li, 2018) with -cx asm20 --secondary=no. Secondly, preliminary phasing was used handcrafted python scripts. For each read, when all the alignment positions were on the AA genome, the read was derived from the AA genome (AA group), on the contrary, it was classified as from the BB genome (BB group). When the alignment position exists in both AA and BB genome, it would be judged as Unknown reads (Unknown group); this type of reads would be phased in the next step. Finally, for Unknown reads, we used PP2PG (Feng et al., 2021) with -ax splice–uf --secondary=no -C5 -O6,24 -B4 –MD of Minimap2 and --maxgap=500 -- mincluster=100 of MUMmer (4.0.0beta2) (Marçais et al., 2018) to evaluate the SNPs site. Finally, reads with SNPs variation sites consistent with the AA genome were classified as AA group, while those with sites consistent with the BB genome were classified as BB group.

#### Genome assembly and phasing

A flow chart of genome assembly approach is shown in Fig. 1b. Briefly, combined the PacBio HiFi reads belonging to AA group and the PacBio HiFi reads that were still defined as unknown after two rounds of phasing, and combined with the Hi-C reads phased to the AA group. Then, using hifiasm (0.15.5-r352) with default settings (Cheng et al., 2022) assemble the AA haplotype genome. Combined the PacBio HiFi reads belonging to BB group and the unknown HiFi reads after two rounds of phasing, B subgenome was assembled with the same process. Finally, three haplotypes (A1, A2 and B) of Plantain and Silk were fully resolved at the chromosomal level. For the initial assemblies, we used Khaper (Zhang et al., 2021) to select primary contigs and filter redundant sequences.

#### Construction of pseudochromosomes

The Hi-C reads were aligned to the contigs using Juicer pipeline. Pseudo-chromosome was constructed with Hi-C data using 3D-DNA pipeline (Dudchenko et al., 2017) with default parameters. The results were polished using the Juicebox Assembly Tools (Durand et al., 2016).

### Genome annotation

#### Repeat annotation

A combined strategy based on homology alignment and *de novo* search was used to identify repeat sequences. Tandem Repeat was extracted using TRF (Benson et al., 1999) by *ab initio* prediction. The homolog prediction commonly used Repbase (Bao et al., 2015) database employing RepeatMasker (Zhi et al., 2006) software and its in-house scripts (RepeatProteinMask) with default parameters to extracted repeat regions. And *ab initio* prediction built *de novo* repetitive elements database by LTR_FINDER (Xu et al., 2007), RepeatScout, RepeatModeler (Flynn et al., 2020) with default parameters, then all repeat sequences with lengths >100 bp and gap ‘N’ less than 5% constituted the raw transposable element (TE) library. A custom library (a combination of Repbase and our *de novo* TE library which was processed by uclust to yield a non-redundant library) was supplied to RepeatMasker for DNA-level repeat identification. On the other hand, EDTA (Ou et al., 2019) was used for prediction, and the results of RepeatMasker and EDTA were merged as the final set of TEs.

#### Protein-coding gene prediction and functional annotation

1. Homolog prediction. Sequences of homologous proteins were downloaded from Ensembl/NCBI/others. Protein sequences were aligned to the genome using TblastN (v2.2.26; E-value ≤ 1e-5) (Altschul et al., 1990), and then the matching proteins were aligned to the homologous genome sequences for accurate spliced alignments with GeneWise (v2.4.1) software (Madeira et al., 2019) which was used to predict gene structure contained in each protein region.
2. *Ab initio* prediction. For gene predication based on *ab initio*, Augustus (v3.2.3), Geneid (v1.4), Genescan (v1.0), GlimmerHMM (v3.04) and SNAP were used in our automated gene prediction pipeline.
3. RNA-seq data. Transcriptome reads assemblies were generated with Trinity (v2.1.1) (Grabherr et al., 2011) for the genome annotation. To optimize the genome annotation, the RNA-Seq reads from different tissues which were aligned to genome fasta using Hisat (v2.0.4) (Kim et al., 2019) / TopHat (v2.0.11) (Trapnell et al., 2009) with default parameters to identify exons region and splice positions. The alignment results were then used as input for Stringtie (v1.3.3) (Pertea et al., 2015) with default parameters for genome-based transcript assembly. The non-redundant reference gene set was generated by merging genes predicted by three methods with EvidenceModeler (EVM, v1.1.1) using PASA (Program to Assemble Spliced Alignment) terminal exon support and including masked transposable elements as input into gene prediction. Finally, Individual families of interest were selected for further manual curation by relevant experts.
4. Functional annotation. Gene functions were assigned according to the best match by aligning the protein sequences to the SwissProt (Bairoch et al., 2000) using Blastp (with a threshold of E-value ≤ 1e-5) (Altschul et al., 1990). The motifs and domains were annotated using InterProScan (v5.31) (Zdobnov et al., 2001) by searching against publicly available databases, including ProDom, PRINTS, Pfam, SMRT, PANTHER and PROSITE. The Gene Ontology (GO) IDs for each gene were assigned according to the corresponding InterPro entry. We predicted proteins function by transferring annotation from the closest BLAST hit (E-value <10^-5^) in the SwissProt database and DIAMOND (v0.8.22) / BLAST hit (E-value <10^-5^) hit (E-value <10^-5^) in the NR database. We also mapped gene set to a KEGG pathway and identified the best match for each gene.

#### Non-coding RNA annotation

The tRNAs were predicted using the program tRNAscan-SE (Chan et al., 2009) (http://lowelab.ucsc.edu/tRNAscan-SE/). For rRNAs are highly conserved, we choose relative species’ rRNA sequence as references, predict rRNA sequences using Blast (Altschul et al., 1990). Other ncRNAs, including miRNAs, snRNAs were identified by searching against the Rfam database with default parameters using the infernal software (Nawrocki et al., 2013).

### Identification of centromeres

Tandem repeats or satellite DNA sequences are commonly found around the centromeres of many plant (and animal) species (Song et al., 2021) and may be classified as ‘centromeric’ or ‘pericentromeric’. However, earlier studies have indicated that Musa lacks a typical centromeric satellite and that its centromeres are instead composed of various types of retrotransposons, especially Ty3/Gypsy-like elements and a LINE-like element named Nani’a (D’Hont et al., 2012; Belser et al., 2021; Hribová et al., 2010). Furthermore, several elements of chromovirus CRM clade, a lineage of Ty3/Gypsy retrotransposons, were found to be restricted to these centromeric regions (D’Hont et al., 2012; Wang et al., 2022). To determine the distribution position of these transposons, we first integrated the annotation results of RepeatMasker, RepeatScount, RepeatModeler, and EDTA to obtain the distribution position of LINE/L1 transposons (RIL code). We then integrated the annotation results of LTRharvest (Ellinghaus et al., 2008), LTR_Finder, and LTR_retriever (Ou et al., 2018) to obtain the distribution positions of Gypsy-like transposons. Finally, we used TEsorter (Zhang et al., 2022) to further classify the LTR transposons obtained above and obtain the distribution location of CRM. The final pericentromeric position was obtained by combining the regions of LINE/L1, CRM, and Gypsy and manually adjusting them. The transposable elements (TEs) were classified following the Wicker et al. classification (Wicker et al., 2007).

### Identification of NLR Genes

To identify NLR genes, we considered those containing at least one NB, a TIR, or a CCR (RPW8) domain. I.e., LRR or CC motifs alone were not sufficiently considered for NLR identification. As a subdivision, we defined TNLs (at least a TIR domain), CNLs (CC+NB domain), RNLs (at least an RPW8 domain), and NLs (at least an NB domain). Canonical architectures contain only NB (Pfam accession PF00931), TIR (PF01582), RPW8 (PF05659), LRR (PF00560, PF07725, PF13306, PF13855) domains, or CC motifs (Van et al., 2019).

### Identification of WRKY Genes

To comprehensively identify WRKY genes, Hidden Markov Model (HMM) seed file of the WRKY domain (PF03106) was obtained from Pfam database (http://pfam.sanger.ac.uk/). HMMER 3.3 (Mistry et al., 2013) was used to search WRKY genes from Plantain and Silk genome database with an E-value threshold of 1e-5. Subsequently, all non-redundant WRKY protein sequences were validated for the presence of WRKY domain by submitting them as search queries to the Pfam and SMART (http://smart.embl.de/) databases. Each potential gene was then manually examined to ensure the conserved heptapeptide sequence at the N-terminal region of the predicted WRKY domain.

### Identification of SNPs, InDels and Structural variation

Plantain and Silk genomes and their haplotypes with each other were aligned using MUMmer with parameters settings ‘-g 1000 -c 90 -l 40’. The alignment block was then filtered out of the mapping noise and the one-to-one alignment was identified by delta-filter with parameters settings ‘-r -q’. Show-snps was used to identify SNPs and InDels (<100 bp) with parameter setting ‘-ClrTH’. The SNPs and InDels were annotated using SnpEff (Cingolani et al., 2012).

To identify inversions and translocations, we aligned the Plantain and Silk genomes and their haplotypes with each other using MUMmer. For the original alignment block to be filtered, we picked a unique alignment block that was longer than 1,000 bp. SyRI (Goel et al., 2019) was used to identify inversions and translocations on both sides. We used the method of (Sun et al., 2018) to identify genes with large structure variations, which mapped gene sequence (including -2 kb upstream and +2 kb downstream of each gene) to query genomes using BWA-MEM (Vasimuddin et al., 2019).

### Identification of PAVs

The potential PAVs in Plantain and Silk genomes and their haplotypes were identified using show-diff in MUMmer (Kurtz et al., 2004). First, sequences that intersected with gap region in the respective genome were excluded. On the other hand, sequence with feature type ‘BRK’ was filtered out, which was considered as non-reference sequence which aligned to the gap-start or gap-end bounder. The gene having >80% overlap with PAV region was considered as a PAV-related gene.

### Identification of HEs

To identify HEs between the haplotypes of Plantain and Silk, we aligned Illumina reads to the DH-Pahang and DH-PKW reference genomes using BWA-MEM and preserved unique alignments. The HE loci were identified based on the depth of read coverage.

### Gene families of eleven bananas in the Musaceae

As references, protein sequences from nine species (*M. textilis* (Abaca), *M. troglodytarum* (Utafun), *M. schizocarpa* (Schizocarpa), *M. acuminata* ssp. *malaccensis* (DH-Pahang), *M. acuminata* ssp. *banksii* (Banksii), *M. acuminata* ssp*. burmannica* (Calcutta 4), *M. acuminata* ssp. *zebrina* (Maia oa), *M. balbisiana* (DH-PKW) and *Ensete glaucum*) were downloaded from phytozome (Goodstein et al., 2012) database. In cases where genes had alternative splicing variants, the longest transcript was selected to represent the gene, and the similarities between sequence pairs were calculated using BlastP (with an E-value cutoff of 1e-10). Furthermore, to identify gene family membership based on overall gene similarity, we employed OrthoMCL (v2.0.9) (Li et al., 2003) with default parameters in conjunction with Markov Chain Clustering.

### Phylogenomic analysis

2043 single-copy orthologous genes were extracted from OrthoFinder (Emms et al., 2019) results and protein sequences were aligned by MAFFT (Katoh et al., 2009). Conserved sites from multiple sequence alignment results were then extracted by Gblocks (Castresana, 2000) and a phylogenetic tree was constructed by RAxML (Stamatakis, 2015) with the *E. glaucum* datasets as the out-group, and 1,000 bootstrap analysis were performed to test the robustness of each branch. Divergence time estimates were calculated by MCMCTree (Puttick, 2019) with two secondary calibration points obtained from previous results, ∼5.4 and ∼9.8 million years ago (mya) for the split time of *M. balbisiana*, *M. acuminata* and *E. glaucum*, *M. acuminata*, respectively. Last, the iTOL (Letunic and Bork, 2021) tools were used to visualize the phylogenetic tree. Gene families undergoing expansion or contraction were identified in the eleven sequenced species using CAFE (p-value thresholdc:=c:0.05, and automatically searched for the λ value) (Han et al., 2013). Genes belonging to significant expanded gene families were subjected to functional analysis by GO and KEGG enrichment.

### Ancestor traceability

While no subspecies has been defined so far in *M. balbisiana*, *M. acuminata* is further divided into multiple subspecies, among which at least four have been identified as contributors (*M. acuminata* ssp. *banksii*, *M. acuminata* ssp. *zebrina*, *M. acuminata* ssp. *burmannica*, and *M. acuminata* ssp. *malaccensis*) to the cultivated banana varieties (Perrier et al., 2011). First, MUMmer (4.0.0beta2) (Kurtz et al., 2004) was used to mapped Plantain and Silk to *M. acuminata* ssp. *banksii*, *M. acuminata* ssp. *Malaccensis, M. acuminata* ssp. *zebrina*, *M. acuminata* ssp. *burmannica*, *M. schizocarpa* and *M. balbisiana* (DH-PKW+PKW). Mapping results were filtered with parameters ‘-i 90 -l 1000’ by delta-filter and the program show-snps was used to identify SNPs between every pair of genomes with parameters ‘-C -T -r -l -x 1’. Set each window with 100, 200, 500 and 1000 kb to divide the genome, use BEDtools (Quinlan, 2014) coverage to count the proportion of SNPs in each window, according to the number of SNPs (for unmatched windows, the number of SNPs is manually set to NA). Using the self-written python script, each window was derived from which ancestors scoring judgment. Secondly, in order to reduce false positives, the SNPs results were adjusted with the collinear block (1 kb, 10 kb and 20 kb) of the genomes. Thirdly, the results were compared with those of previous studies (accession numbers: Plantain 148 and 149, Silk 139 and 140) (Martin et al., 2023). Combining the above steps, the final results were obtained.

### Analysis of synteny and whole-genome duplication

Syntenic blocks were identified using jcvi (MCScanX (Wang et al., 2012) Python version with default parameters. Proteins were used as queries in searching against genomes of other plant species to find the best matching pairs. Each aligned block represented an orthologous pair derived from the common ancestor. In general, the ratio of nonsynonymous substitution rate (Ka) and synonymous substitution rate (Ks) was used to assess gene selection by PAML. As input files, the sequences of the homologous genes were imported into WGDI (Sun et al., 2021) to calculate the gene pair values.

### Statistics of lost homologous gene pairs

We aligned DH-Pahang, DH-PKW, and sub-genomes of Plantain and Silk using GeneTribe (Chen et al., 2020), respectively, and examined the presence/absence of orthologous pairs in the Pa/Pb and Sa/Sb genomes. We extracted 1:1 ortholog pairs shared by DH-Pahang and DH-PKW and examined the presence/absence of orthologous pairs in the Pa/Sa and Pb/Sb genomes. We then selected a subset of genes that had lost their orthologous pair either in Pa/Pb or Sa/Sb genomes but not in all genomes to study the mechanism of gene fractionation. A genome-wide Chi-squared test was performed to determine whether the number of DH-Pahang/Pa lost genes differed significantly from the number of DH-PKW/Pb lost genes (P ≤ 0.05). Genes in each orthologous pair were categorized as singletons or duplicates based on their duplication status in DH-Pahang and DH-PKW genomes. For orthologous pairs where one gene was annotated as a singleton and another as a duplicate, we calculated the percentage of lost genes in each category and tested for deviation from a 1:1 ratio using a Chi-squared test. We measured the Ka/Ks values for lost and conserved orthologous gene pairs based on their counterparts from DH-Pahang and DH-PKW genomes. We used the sequences of lost genes from *M. acuminata* or *M. balbisiana* genomes as queries and mapped them back to the Plantain and Silk genomes using BLASTN v2.7.1 (evalue 1e–10; word_size 30; -qcov_hsp_perc 0.8) (Altschul et al., 1990) to study the segmentation/deletion mechanism.

### Statistics of expression bias of homologous genes

The triallelic data obtained by MCscan (Tang et al., 2008) and the expression result file obtained by RSEM (Li et al., 2011) were used to obtain the corresponding expression levels. The expression level of sub-genome A was calculated as (A1+A2)/2, and expressed genes with TPM>=1 were selected as candidate genes. A Chi-square test was then performed to determine whether the expression of sub-genome A significantly differed from that of sub-genome B (P ≤ 0.05), thus identifying the sub-genome with dominant expression. To examine the effect of TE insertion on gene expression, the distance of the nearest TE inserted into the upstream region of a gene was identified using BEDtools (Quinlan, 2014) (closest -id -D a), and the correlation was compared for the orthologous pairs between the A and B genomes.

### Identification of alleles

To identify the homologous regions between three haplotypes of Plantain and Silk, we applied the MCscan with lastal parameters ‘--cscore=.99’ to contain reciprocal best hit (RBH) for construct the syntenic blocks based on well-aligned genes. Allelic gene pairs were selected according to the following rules: (1) paired regions must be on homologous haplotypes; (2) when there is one-to-many paired genes, take the one with the higher C-score (score(A, B) / max(score(A,), score(,B))); (3) the three genes are paired with each other are identified as 3 alleles, the two genes are paired with each other are identified as 2 alleles, the others are 1 allele; (4) Syntenic gene pairs defined above were double-checked manually.

### Identification of differential expression allelic genes (DEA)

RNA samples from 35 Plantain and 23 Silk sets were trimmed using the Trimmomatic program (Bolger et al., 2014) and mapped against annotated gene models using STAR/2.7.3a (Dobin et al., 2013), with only the best alignment retained for each read using the parameters --twopassMode Basic --outSAMmultNmax 1. The RSEM program (Li et al., 2011) was then used to estimate TPM values.

1. RNA data without duplicates. First, sort TPM from high to low (I, II, III) of 3 alleles, second, identify DEA adopted five standards:

1. TPM_A1_ >= 1 or TPM_A2_ >= 1 or TPM_B_ >= 1;
2. Count_A1_ >=10 or Count_A2_ >=10 or Count_B_ >= 10;
3. I/II or II/III, more than twofold difference (Alleles 3);
4. TPM_A1_ / TPM_A2_ >= 2 or <= 0.5; TPM_A1_ / TPM_B_ >= 2 or <= 0.5; TPM_A2_ / TPM_B_ >= 2 or <= 0.5 (Alleles 2);
5. Detected in at least 2 samples.
2. RNA data with biological replicates, DEA was determined if the log fold change of TPM values between two alleles was greater than 2 with adjusted P value < 0.05, and was detected in at least 2 samples.

### Culture of Foc-TR4 strain and preparation of inoculant

The strains were provided by the Guangdong Provincial Key Laboratory of Tropical and Subtropical Fruit Tree Research. The 1 cm^2^ size bacteria were taken in the ultra-clean workbench and inoculated into 50 ml of sterile Potato Dextrose Broth (PDB) medium, 28 c: for about 7 days. Aspirate the spore suspension, observe the number of spores under a light microscope using a hemocytometer, and measure the concentration of the spore suspension to be 1×108 ml^-1^. Dilute the spore suspension to 1×106 ml^-1^ with sterile water for use.

### Inoculation and resistance evaluation (potted and field evaluation)

Banana seedlings that have grown to 4-6 leaves are removed from the substrate, rinsed, and their roots completely immersed in the spore suspension for 30 minutes. Using the “double-pot system” of (Mohamed et al., 2001), the seedlings were planted in a nutrient cup (20cm in diameter, 15cm in bottom diameter, 17.5cm in height) containing sterilized perlite, and the nutrient cup was placed in a 50cm long, in a plastic box with a width of 30cm and a height of 10cm without a lid, there is tap water at the bottom of the box, the water depth is 1-2cm, and the hoagland nutrient solution is regularly poured. Set Silk as the susceptible control and Plantain as the resistant control. At the time of uninoculation (week 0) and the first to fifth weeks after inoculation, 3 plants of each variety were taken from the inoculated treatment group and the non-inoculated control group (only the control group was take^n^ in the 0th week), and the whole plant was longitudinally cut. Photographs were taken and the rhizomes were taken as samples, and the samples were stored at −80°C. Referring to the method of (Viljoen et al., 2017), the rhizome discoloration index (RDI) was calculated according to the discoloration inside the rhizome to evaluate the disease resistance of bananas, 5<RDI≤6 is high sensitivity.

The field evaluation was carried out in the experimental field of Banana and Vegetable Research Institute in Dongguan City, Guangdong Province, where the soil had been infected with Foc-TR4, and the incidence rate of ’Cavendish’ planted in this field was not less than 70% (Zuo et al., 2018). No chemicals were applied during the test. Two evaluations were conducted in November 2020 and November 2021. When the plant became ill or the test was over, the rhizome and pseudostems were cut and photographed.

### Analysis of Plantain and Silk resistance to Foc-TR4 in five stages using RNA seq

RNA-seq was performed on three biological replicates of post-inoculated and uninoculated rhizomes at five different developmental stages (1-5 weeks) of Plantain and Silk. Trimmomatic (Bolger et al., 2014) was used to remove low-quality reads, and clean reads were then mapped to the reference genomes of Plantain and Silk using STAR/2.7.3a (Dobin et al., 2013). The mapping reads corresponding to each transcript were assembled, and TPM values were calculated using RSEM. DEG analysis was conducted using DESeq2 from the R Bioconductor package, with log2FC > 2 for genes with increased transcript abundance and log2FC < −2 for genes with decreased transcript abundance, and a P-value threshold of ≤ 0.05. Comparisons were made between post-inoculation and control groups at weeks 1 to 5 for both Plantain and Silk.

### Co-expression network between differentially expressed MYB transcription factors and lignin biosynthesis genes after inoculation

The co-expression algorithm in R package WGCNA (Langfelder et al., 2008) was used to identify co-expression modules. The power value threshold option was disabled while constructing modules, and the obtained power values ranged from 1 to 20. To determine the average and independence connection degrees of multiple modules, the gradient technique was employed. A degree of independence of 0.8 was considered suitable for the power value. Modules were built using the WGCNA method once the power value threshold was established, and genes related to each module were examined. To ensure the findings’ high reliability, the minimum number of genes in a module was set at 30. Co-expression networks were visualized using Cytoscape (Shannon et al., 2003). MYB TFs typically recognize specific AC-rich cis elements ([ACC(A/T)A(A/C)(T/C)]) that are especially prevalent in the promoters of PAL, 4CL, CCR, and CAD (Zhao et al., 2011), regulating the lignin biosynthesis.

## DATA AVAILABILITY

The Plantain and Silk genome assembly have been deposited in the Genome Warehouse in National Genomics Data Center, Beijing Institute of Genomics, Chinese Academy of Sciences / China National Center for Bioinformation, under project number PRJCA015888 that is publicly accessible at https://ngdc.cncb.ac.cn/gwh.

## AUTHOR CONTRIBUTIONS

L.-L.C., O.S., J.-M.S. and G.Y. conceived and supervised this study. O.S., G.Y., W. H., F. B, Y. L, T. D., G. M., S. L., C. L., Q. Y, C. H., H. G., and T. D. collected samples and performed experiments. W.-Z.X., Y.-Y.Z., L.-L.C., J.-M.S., R.Z., Y.-X. G., W.-H. Z., M.-H. Y., S.-J. P., X.-T. Z., X.-D. X., Z.-W. Z., J.-W. F. and J. Z. performed genome assembling and annotation, comparative genomics analysis, and transcriptome data analysis. J.D. and Q.G. performed karyotype analysis of Plantain and Silk banana, W.-Z.X., Y.-Y.Z., J.-M.S., O.S. and L.-L.C. wrote and revised the paper.

## ACKNOWLEDGEMENTS

We sincerely thank professor Maojun Wang in Huazhong Agricultural University for his guidance in asymmetric evolution of Plantain and Silk genomes. We also acknowledge the computing platform of the National Key Laboratory of Crop Genetic Improvement in HZAU for providing the computational resources.

## ONLINE CONTENT

Any methods, additional references, research reporting summaries, source data, statements of code and data availability and associated accession codes are available online.

